# The RNA binding protein Swm is critical for *Drosophila melanogaster* intestinal progenitor cell maintenance

**DOI:** 10.1101/2022.01.03.474858

**Authors:** Ishara S. Ariyapala, Kasun Buddika, Heather A. Hundley, Brian R. Calvi, Nicholas S. Sokol

## Abstract

The regulation of stem cell survival, self-renewal, and differentiation is critical for the maintenance of tissue homeostasis. Although the involvement of signaling pathways and transcriptional control mechanisms in stem cell regulation have been extensively investigated, the role of post-transcriptional control is still poorly understood. Here we show that the nuclear activity of the RNA-binding protein Second Mitotic Wave Missing (Swm) is critical for *Drosophila* intestinal stem cells (ISCs) and their daughter cells, enteroblasts (EBs), to maintain their identity and function. Loss of *swm* in these intestinal progenitor cells leads ISCs and EBs to lose defined cell identities, fail to proliferate, and detach from the basement membrane, resulting in severe progenitor cell loss. *swm* loss further causes nuclear accumulation of poly(A)+ RNA in progenitor cells. Swm associates with transcripts involved in epithelial cell maintenance and adhesion, and the loss of *swm*, while not generally affecting the levels of these Swm-bound mRNAs, leads to elevated expression of proteins encoded by some of them, including the fly orthologs of Filamin and Talin. Taken together, this study indicates a role for Swm in adult stem cell maintenance, and raises the possibility that nuclear post-transcriptional gene regulation plays vital roles in controlling adult stem cell maintenance and function.

## INTRODUCTION

Understanding the biology and behavior of stem cells and their niches is important because of their critical role in maintaining adult tissue homeostasis throughout life. Stem cell proliferation, survival, and differentiation are tightly regulated by both cell-autonomous and non-cell-autonomous factors whose mis-regulation can compromise tissue integrity (Losick et al., 2011; Morrison and Spradling, 2008). Despite the interest in studying stem cell behaviors, how stem cell populations maintain cell identity and stemness to deliver proper cell and tissue functions is less well understood.

The *Drosophila* midgut epithelium serves as an ideal model to study stem cell behaviors because this high turnover tissue hosts a stem cell population, known as intestinal stem cells (ISCs), that gives rise to a simple cell lineage (fig. 1A). In this lineage, intestinal stem cells (ISCs) are basally located and physically attached to a basement membrane that neighbors a layer of visceral muscle. ISCs are the only mitotic cells in the midgut and often divide asymmetrically to produce a new ISC and a transient enteroblast (EB) that differentiates into an enterocyte (EC) through activation of the Notch signaling pathway (Biteau et al., 2011; Micchelli and Perrimon, 2005; Ohlstein and Spradling, 2005; Ohlstein and Spradling, 2007). At the same time, a small percentage of ISCs generate enteroendocrine (ee) cells through asymmetric division or direct differentiation (Amcheslavsky et al., 2014; Biteau and Jasper, 2014; Guo and Ohlstein, 2015; Zeng and Hou, 2015). ISCs and EBs are collectively known as intestinal progenitors. Because the transcriptomes of the two cell types are largely similar, there are multiple markers that identify progenitors but few that are specific to either ISCs or EBs (Dutta et al., 2015; Hung et al., 2020). Progenitors express the transcription factor Escargot (Esg) while ISCs are identified by the expression of the Notch ligand, Delta (Dl), and EBs are identified as cells that express Notch signaling reporters. Ee cells express the transcription factor Prospero (Pros) and ECs express the transcription factor Pou domain-1 (Pdm-1) (fig. 1A) (Jiang et al., 2011; Micchelli and Perrimon, 2005; Ohlstein and Spradling, 2005). Unlike many other stem cell populations, ISCs are not associated with any specialized supporting cells making it difficult to define their exact niche. Growing evidence, however, supports the notion that the visceral muscle and basement membrane, including its extracellular matrix (ECM) proteins, serve as a basal niche to support ISCs (Biteau and Jasper, 2011; Buchon et al., 2010; Jiang et al., 2009; Lin et al., 2008; O’Brien et al., 2011). A number of signaling pathways, including Wnt, Hippo, Notch, EGFR, Insulin, BMP/Dpp and JAK/STAT, as well as environmental signals, including chemicals, nutrients and pathogens, are involved in regulating ISC proliferation, survival and differentiation as well as maintenance of other cell types (Miguel-Aliaga et al., 2018). Recent studies have indicated that, in addition to the transcriptional regulation provided by these pathways, post-transcriptional regulation also plays critical roles in ISCs. For example, post-transcriptional gene regulation through RNA binding proteins (RBPs) and small RNAs, such as microRNAs (miRNAs), have been shown to add important layers to regulate ISC behaviors both under homeostatic and externally challenged conditions (Buddika et al., 2020; Buddika et al., 2021b; Chen et al., 2015; Foronda et al., 2014; Luhur et al., 2017; Mukherjee et al., 2021; Shanahan et al., 2020). In other stem cell types, such as human and mouse embryonic stem cells as well as *Drosophila* S2R+ cells, nuclear post-transcriptional processes including alternative splicing, polyadenylation, RNA export, and nuclear RNA decay have been shown to regulate potency, self-renewal, and differentiation (Chen and Hu, 2017; Herold et al., 2003; Hurt et al., 2009; Silla et al., 2020; Wang et al., 2013). However, the role of nuclear post-transcriptional processes in regulating stem cell behaviors of *in vivo* tissue based stem cell models remain poorly understood.

**Figure 1.**
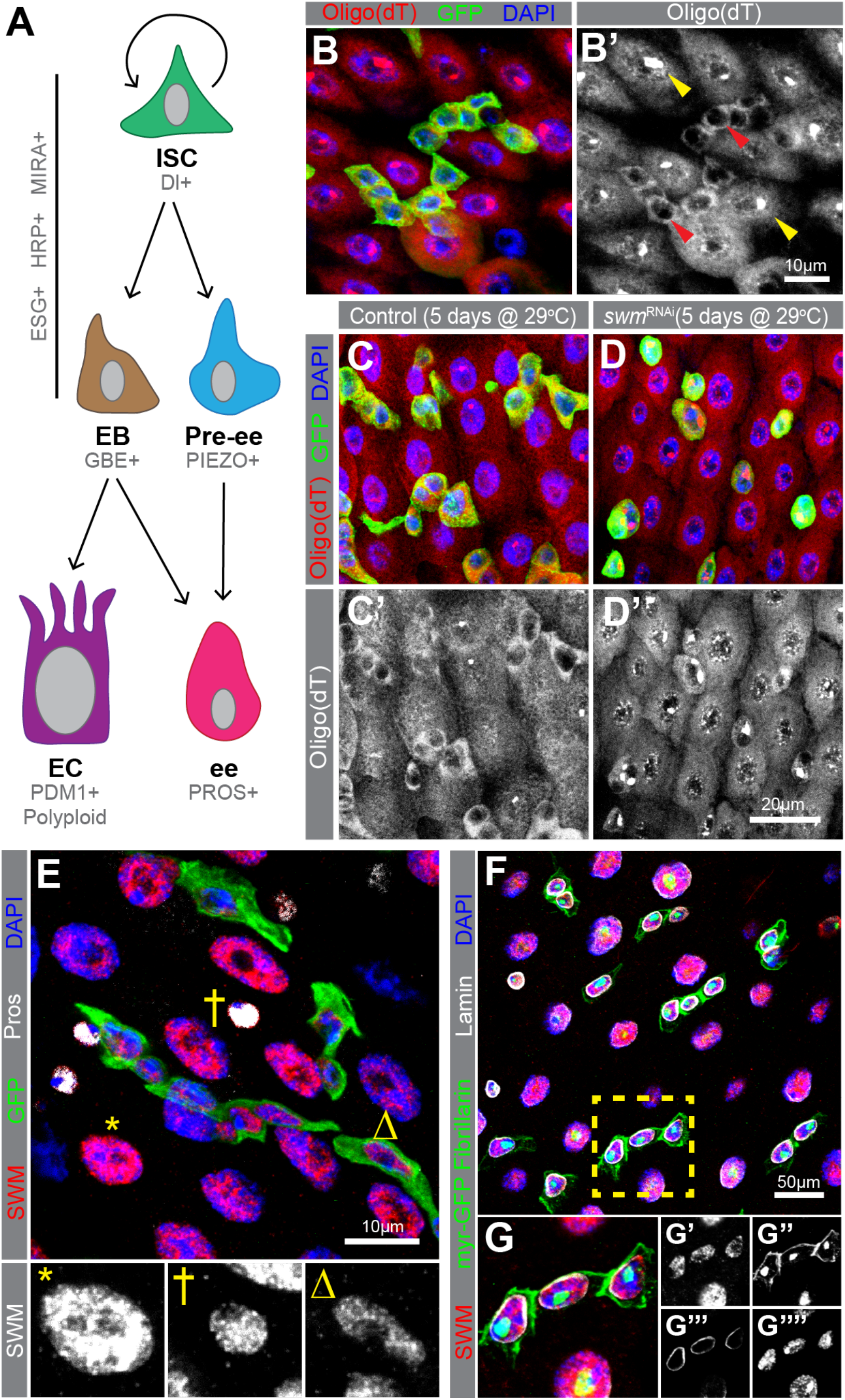
Knockdown of *swm* in intestinal progenitor cells leads poly(A)+ RNA accumulation in nuclei. (A) A schematic of intestinal cell types and markers. (B) Poly(A)+ RNA distribution in cells from *esg*^TS^ intestines, labelled by oligo (dT) probes (red) and Ps are labelled with GFP (anti-GFP in green) and nuclei with DAPI (blue). (B’) Poly(A)+ RNA distribution is shown in gray scale in Ps (red arrow heads) and ECs (yellow arrow heads). (C and D) Sections from (C) *esg*^TS^ and (D) *esg*^TS^; *swm* ^RNAi^ PMGs after 5 days at 29°C stained with oligo (dT) (red), GFP (green), and DAPI (blue). (E) PMG section from *esg*^TS^ stained for Swm (anti-Swm in red). Ps are shown in GFP (green), ees are stained for Prospero (anti-Prospero in white) and nuclei for DAPI (blue). Enlarged EC (*), ee (†) and P (Δ) show Swm in gray scale. Individual channels of panel E are shown in Figure S1. (F and G) PMG region from *UAS-myr GFP* driven by *esg*-*GAL4* stained for Swm (anti-Swm in red) and counterstained for cell membrane of Ps (*myr-GFP* in green), nucleoli of all cells (anti-Fibrillarin, nuclear green), nuclear membranes of all cells (anti-Lamin in white) and nuclei (DAPI in blue). (G) Enlargement of cells indicated in E with individual channels for Swm in red (G’), cell membranes (*myr-GFP*) and nucleoli (Fibrillarin) in green (G’’), nuclear membranes in white (G’’’) and nuclei in blue (G’’’’). Complete genotypes are listed in table S1. P, progenitor cell; EC, enterocyte; ee, enteroendocrine cell; Pros, Prospero; PMG, posterior midgut.

Here, we used the *Drosophila* midgut as a tissue-based stem cell model to study nuclear post-transcriptional regulatory mechanisms. We identify the RBP Second Mitotic Wave Missing (Swm) as a critical factor in maintaining intestinal progenitors and post-transcriptionally regulating mRNAs. Our study reveals the presence and the critical importance of nuclear RBP activity for *Drosophila* intestinal progenitor cell survival and maintenance.

## MATERIALS AND METHODS

### *Drosophila* strains and husbandry

Age matched mated female flies were used in all experiments. All fly strains were cultured on standard Bloomington media (https://bdsc.indiana.edu/information/recipes/bloomfood.html) in 18°C, 25°C or 29°C incubators set for a 12hr light/dark schedule and 65% humidity. The full genotypes of all fly strains used in this study are listed in Table S1.

### Temperature

For temporal and regional gene expression-targeting (TARGET) experiments, flies were reared at 18°C, upon eclosion collected progeny over 2 days, aged 2 more days at 18°C and shifted to 29°C up to 10 days before being dissected. For clonal analysis using mosaic analysis with repressible cell marker (MARCM) technique, flies were grown at 25°C until eclosion, collected 0-2 days old animals and heat-shocked immediately at 38°C for 40min in a Lauda circulating water bath. Following heat-shock, flies were reared at 25°C up to 10 days.

### Construction of new strains

*swm*^RNAi-2^ was generated by first predicting effective shRNAs targeting the swm 3’UTR using splashRNA (Pelossof et al., 2017) and then annealing and subcloning oligos encoding one of these shRNAs (TTAACAATTATATATCCGCGTA) into the EcoRI and NheI sites of pWalium20. The transgene was subsequently inserted into the *VK33* landing site by Rainbow Genetics (Camarillo, CA).

### Antibody generation

Anti-Swm antibodies were generated in rats (Cocalico Biologicals) against a 6×HIS-tagged fragment of the first 220 amino acids of Swm that was expressed and purified according to standard methods. This Swm-encoding plasmid (pNIK1384) was generated by PCR amplifying the *swm* coding region from cDNA LD45403 (obtained from the *Drosophila* Genomics Resource Center using oligonucleotides 4290 (acgaccgaaaacctgtattttcagggcgccATTCTGGAGAATTCGGACAAGCTCAAGGAT), 4292 (actagttgagctcgtcgacgtaggcctttgTCAAAGACCTGCTCCGCCGGGACCACCTCC), and high-fidelity Q5 polymerase (NEB), subcloning the resulting PCR product into the NcoI and EcoRI sites of pHIS.parallel (Sheffield et al., 1999) using HiFi DNA assembly master mix (NEB), and sequence verifying the resulting plasmid to confirm the absence of any PCR-induced errors.

### Dissections and immunostaining

Gastrointestinal tracts of adult female flies were dissected in 1x PBS (137mM NaCl, 2.7mM KCl, 10mM Na_2_HPO_4_, KH_2_PO_4_, pH 7.4), fixed in 4% paraformaldehyde (Electron Microscopy Sciences, Cat. No. 15714) in 1x PBS for 45 min, washed three times in 1xPBT (1xPBS, 0.1% Triton X-100) and then blocked (1x PBT, 0.5% bovine serum albumin) for 45min. Subsequently, samples were incubated at 4°C overnight with primary antibodies, including mouse anti-Prospero (MR1A, Developmental Studies Hybridoma Bank, 1:100), mouse anti-V5 (MCA1360GA, Bio-Rad, 1:250), and rabbit anti-GFP (A11122, Life Technologies, 1:1000), rat anti-Swm (this study, 1:50), rat anti-Cheerio (Sokol and Cooley, 1999), (1:1000), mouse anti-Talin (A22A and E16B, Developmental Studies Hybridoma Bank, 1:1 mixture, 1:50 each), mouse anti-Lamin (ADL67.10, Developmental Studies Hybridoma Bank, 1:50), mouse anti-Coracle (C566.9, Developmental Studies Hybridoma Bank, 1:50), mouse anti-Discs large (4F3, Developmental Studies Hybridoma Bank, 1:100), mouse anti-Headcase (U33, Developmental Studies Hybridoma Bank, 1:3), mouse anti-Osa (Osa, Developmental Studies Hybridoma Bank, 1:20), mouse anti-Shot (mAbRod1, Developmental Studies Hybridoma Bank, 1:20), and rabbit anti-Fibrillarin (ab5821, Abcam, 1:500). The following day, samples were washed in 1xPBT and incubated for 2–3 hours with secondary antibodies, including AlexaFluor 488- and 568-conjugated goat anti-rabbit, -mouse, -rat and -chicken antibodies (Life Technologies, 1:1000). AlexaFluor 647-conjugated goat-HRP antibodies were used in the secondary antibody solution whenever required.

Finally, samples were washed multiple times in 1xPBT, including one wash with 5μg/ml DAPI in PBT, and mounted in Vectashield mounting medium (Vector Laboratories). An alternative staining protocol was used for mouse anti-Delta (C594.9B, Developmental Studies Hybridoma Bank, 1:500) staining as described in (Buddika et al., 2021a) and these samples were mounted in ProLong Diamond mounting medium (Invitrogen, P36970). Intestines stained with anti-Delta antibody were also stained with anti-GFP antibody, since the methanol steps required in this protocol quenched GFP fluorescence. Cell death analysis was performed using the ApopTag® Fluorescein *in situ* Apoptosis Detection Kit (Sigma-Aldrich, Cat. No. S7110) following manufacturer’s instructions.

### Microscopy and image processing

Images of whole dissected intestines were collected on a Zeiss Axio Zoom microscope. Images of immunostained intestines were collected on a Leica SP8 Scanning Confocal microscope. Samples to be compared were collected under identical settings on the same day, image files were adjusted simultaneously using Adobe Photoshop CC, and figures were assembled using Adobe Illustrator CC. Image J FIJI (https://fiji.sc/) was used to quantify the fluorescence intensity.

### Oligo-dT fluorescent *in situ* hybridization

Oligo-dT fluorescent *in situ* hybridization was performed as described in Buddika et al., 2020 and briefly, adult female intestines were dissected out in ice cold 1X PBS and fixed in 4% w/v paraformaldehyde (Electron Microscopy Sciences, Cat. No. 15714) in PBS for 45 min while rocking. Tissue was then washed in 1xPBT (1xPBS, 0.3% v/v Triton X-100) 3 times 5 min each. When protein immunostaining is required, samples were blocked in RNase free blocking solution (0.3% v/v PBT, 0.5% ultra-pure Bovine Serum Albumin) (Ambion, cat. no. AM2616) for 45 min and primary and secondary antibody stainings were carried out. After secondary antibody staining, samples were washed 3x in 0.3% v/v PBT and subjected to a second sample fixation in 4% w/v paraformaldehyde (Electron Microscopy Sciences, Cat. No. 15714) in PBS for 45 min while rocking. Tissue was then washed in 1xPBT (1xPBS, 0.3% v/v Triton X-100) 3 times 5 min each. Subsequently, samples were gradually dehydrated in a series of 0.3% v/v PBT (1xPBS, 0.3% v/v Triton X-100): Methanol (7:3, 1:1, 3:7) washes and incubated in 100% Methanol for 10 min. Next, tissue was rehydrated with a series of 0.3% v/v PBT: Methanol (3:7, 1:1, 7:3) washes and finally washed in 0.3% v/v PBT for another 5 min. Then samples were rinsed once in hybridization wash buffer (20% formamide, 2x SSC, DEPC-treated water) and washed in hybridization wash buffer for 10 min. Then the wash buffer was completely removed and oligo-dT probes were added in 50μl of ULTRAhyb^TM^-Oligo buffer (Ambion, cat. no. AM8663). Samples were incubated overnight at 37°C. The following day, probe solution was removed, and samples were washed in hybridization wash buffer three times for 20 min each and DAPI was added in the second wash. Finally, samples were mounted using ProLong Diamond mounting solution (Invitrogen, Cat. No. P36971).

### RNAscope *in situ* hybridization

RNAscope *in situ* hybridization was performed as described in Buddika et al., 2021b with some changes. Briefly, adult female flies were dissected in ice cold 1X PBS and fixed in 4% w/v paraformaldehyde (Electron Microscopy Sciences, Cat. No. 15714) in PBS for 2 hours while rocking. Tissue was then washed in 1xPBT (1xPBS, 0.3% v/v Triton X-100) 3 times 5 min each. Samples were gradually dehydrated in a series of 0.3% v/v PBT (1xPBS, 0.3% v/v Triton X-100): Methanol (7:3, 1:1, 3:7) washes and incubated in 100% Methanol for 10 min. Then tissue was rehydrated with a series of 0.3% v/v PBT: Methanol (3:7, 1:1, 7:3) washes and finally washed with 0.3% v/v PBT for another 5 min. For next steps, reagents from RNAscope Multiplex Fluorescent Reagent Kit v2 assay were used. Fixed tissue was transferred into 0.2 mL PCR tubes and incubated in RNAscope Protease III reagent for 5 min at 40°C (all the incubations at 40°C were done in a PCR thermal cycler). Samples were immediately washed with 1X PBS twice. RNAscope probes for *cher* (50X) (pre warmed to 40°C) (ACD Bio., Cat. No. 1050721-C2) was diluted in RNAscope Probe Diluent (ACD Bio., Cat. No. 300041) to have 1X concentration and added 20μl per sample. Next, samples were incubated at 40°C overnight. Following day, samples were washed twice with 1X RNAscope wash buffer. Next, RNAscope Multiplex FL v2 AMP 1, RNAscope Multiplex FL v2 AMP 2, RNAscope Multiplex FL v2 AMP 3 and RNAscope Multiplex FL v2 HRP-C2 steps were done as described in Chapter 4 of Fluorescent v2 assay manual, ACD Bio. Finally, samples were incubated for 30 min at 40°C with Opal 620 (AKOYA Biosciences, Cat. No. FP1495001KT, 1:1500 in TSA buffer) and washed with 1X RNAscope wash buffer and counterstained with DAPI. Samples were mounted in ProLong Diamond mounting medium (Invitrogen).

### Bleomycin feeding assay

Flies were reared on standard Bloomington *Drosophila* stock center media for an appropriate time at 18°C or 29°C and divided into two groups prior to the assay. Group 1 and 2 were transferred into vials containing a chromatography paper soaked in 5% sucrose in water (control) and 5% sucrose and 25µg/ml bleomycin in water respectively. Flies were shifted to 18°C or 29°C incubators based on the experiment and dissected after 24 hours of feeding.

### Western blot analysis

Adult female flies were used for protein isolation. Whole adult flies were lysed in I-RIPA protein lysis buffer (150mM NaCl, 50mM Tris-HCl pH 7.5, 1mM EDTA, 1% Triton X-100, 1% Na Deoxycholic Acid, 1xprotease inhibitor cocktail). Prepared protein extracts were resolved on a 4-20% gradient polyacrylamide gel (Bio-Rad, Cat. No. 456-1093), transferred to Immobilon^®^-P membrane (Millipore, Cat. No. IPVH00010) and probed with rat anti-Swm (this study,1:100), rabbit anti-GFP (ab290, Abcam, 1:10,000) or mouse anti-a-tubulin (12G10, Developmental Studies Hybridoma Bank, 1:1000) antibodies. After washing with 1xTBST (1xTBS, 0.1% Tween-20) blots were incubated with anti-rat, -rabbit, -or -mouse conjugated HRP secondary antibodies. Subsequently, blots were washed with 1xTBST, treated with ECL-detection reagents (Thermo Scientific, Cat. No. 1859701 and 1859698) and finally exposed to chemiluminescence films (GE Healthcare, Cat. No. 28906839).

### RNA immunoprecipitation, library preparation and qPCR

Gastrointestinal (GI) tracts were dissected from 200 adult female flies (per replicate) in ice-cold 1xPBS. Dissected intestines were placed on a petri dish with minimum amount of 1xPBS as a monolayer and irradiated three times at 2000J in a UV cross-linker, with mixing between each irradiation to maximize surface exposure. Immediately after irradiation, samples were snap frozen in liquid nitrogen and stored at −80°C until all the samples are prepared. Same day, each frozen tissue sample was lysed in 1ml of I-RIPA protein lysis buffer (150mM NaCl, 50mM Tris-HCl pH 7.5, 1mM EDTA, 1% Triton X-100, 1% Na Deoxycholic Acid, 1xprotease inhibitor cocktail, RNase inhibitor). After centrifugation 50µl of supernatant was saved as the total input and the rest was incubated with Swm antibody coated Dynabeads^TM^ Protein G (Invitrogen, Cat. No. 10003D) for 2 hours at 4°C while rocking. Beads were collected on a magnetic stand and washed with fresh lysis buffer containing RNase inhibitor. Subsequently, Swm bound RNA was released from beads using Proteinase K (Ambion, Cat. No. AM2546) treatment and TRIzol® LS reagent (Ambion, Cat. No. 10296028) was used to isolate immunoprecipitated RNA. The rRNA-depleted libraries were prepared using the Ovation® SoLo RNA-seq system (Part No. 0502 including Parts 0407 and S02240) following manufacturer’s instructions. The quality and quantity of final libraries were assessed using Agilent 2200 TapeStation and KAPA Library Quantification Kit, respectively. For qPCR, isolated RNA was first treated with Turbo DNase (ThermoFisher, Cat. No. AM2239) and gDNA-free RNA was used for cDNA synthesis with Superscript III (ThermoFisher, Cat. No. 56575). qPCR was performed using the PowerUp SYBR Green Master Mix (ThermoFisher, Cat. No. A25742) in a StepOnePlus machine (ThermoFisher). Primers for all targets detected are listed in table S5. Transcript levels were quantified in triplicates and normalized to *Gapdh1*. Fold enrichment was calculated as the ratio of transcript in Swm IP versus total input.

### FACS isolation of progenitor cells and RNA-seq library preparation

Gastrointestinal (GI) tracts were dissected from 80-100 adult female flies (per replicate) in ice-cold 1xPBS. Then cells were dissociated by treating intestines with 1mg/ml elastase at 27°C for 1 hour with agitation. Subsequently, ∼25,000-50,000 GFP^+^ intestinal progenitor cells were sorted using a BD FACSAria™ II flow cytometer equipped with a 100µm nozzle at the IUB Flow Cytometry Core Facility. Total RNA was prepared using the TRI reagent (Molecular Research Center, Cat. No. TR118). The rRNA-depleted libraries were prepared using the Ovation® SoLo RNA-seq system (Part No. 0502 including Parts 0407 and S02240) following manufacturer’s instructions. The quality of final libraries was assessed using Agilent 2200 TapeStation and libraries were quantified using KAPA Library Quantification Kit.

### CLIP-seq and RNA-seq data analysis

Swm CLIP and transcriptomic data analysis were performed as described in Buddika et al., 2021a using a python based in-house pipeline (https://github.com/jkkbuddika/RNA-Seq-Data-Analyzer). First, the quality of raw sequencing files was assessed using FastQC (Andrews, 2010) version 0.11.9. Then, reads with low quality were removed using Cutadapt (Martin, 2011) version 2.9. Next, the remaining reads were mapped to the Berkeley *Drosophila* Genome Project (BDGP) assembly release 6.28 (Ensembl release 100) reference genome using STAR genome aligner (Dobin et al., 2013) version 2.7.3a and duplicated reads were eliminated using SAMtools (Li et al., 2009) version 1.10. Subsequently, the Subread (Liao et al., 2019) version 2.0.0 function *featureCounts* was used to count the number of aligned reads to the nearest overlapping feature. All subsequent analysis and data visualization steps were performed using custom scripts written using R. Next, differential gene expression analysis was performed with the Bioconductor package DESeq2 (https://bioconductor.org/packages/release/bioc/html/DESeq2.html) (Love et al., 2014) version 1.26.0. Unless otherwise noted, significantly upregulated and downregulated genes were defined as FDR < 0.05; Log_2_ fold change > 1 and FDR < 0.05; Log_2_ fold change < −1, respectively and were used to identify enriched Gene Ontology (GO) terms using gProfiler2. A selected significantly enriched GO categories were plotted.

## Statistical analysis

For all statistical analyses, GraphPad Prism, Version 9.0 was used. First, D’Agostino-Pearson test was used to test the normality of datasets. For comparisons involved in two datasets, if datasets follow [1] a parametric distribution, an Unpaired t-test or [2] a non-parametric distribution, a Mann-Whitney test was performed. Three or more datasets following a parametric distribution were analyzed using an ordinary one-way ANOVA test. Multiple comparisons of three or more datasets following a non-parametric distribution were analyzed using Kruskal-Wallis test. Unless otherwise noted, significance is indicated as follows: n.s., not significant; *p < 0.05; **p < 0.01; ***p < 0.001; ****p < 0.0001.

## RESULTS

### Nuclear Poly(A)+ RNA accumulates in the absence of *swm*

Previous analyses of RBPs in progenitor cells have focused on their cytoplasmic roles, leaving a possible role for RBPs in progenitor nuclei unexplored (Buddika et al., 2020; Buddika et al., 2021b; Chen et al., 2015; Luhur et al., 2017). To begin investigating this possibility, we first assessed the poly(A)+ RNA distribution in *Drosophila* intestinal cells by oligo d(T) *in situ* hybridization. While oligo d(T) signal was detected in the cytoplasm of all cell types as well as the nuclei of most ECs, it was noticeably absent from the nuclei of progenitor cells (fig. 1B-B’, S1A). This prompted us to ask whether the regulation of RNA localization differed between progenitors and their daughter cells. To address this question, we took a candidate genetic approach and knocked down selected RNA processing and nuclear export related factors in intestinal progenitor cells via RNA interference (RNAi) and observed progenitor cell numbers as well as poly(A)+ RNA distribution detected by oligo d(T) *in situ* hybridization. For our screen, we focused on genes implicated in RNA processing and nuclear export and generated a candidate gene list using the gene ontology (GO) term analysis function on Flybase (Larkin et al., 2021). From the candidate gene list, we selected 43 genes, encoding both general RNA processing and nuclear export factors as well as proteins implicated or predicted to have nuclear RNA processing functions and that were present within the Bloomington *Drosophila* Stock Center RNAi strain collection (see Table S2 for the screen summary). Each RNAi strain was crossed to a strain harboring both the intestinal progenitor cell driver *esg-GAL4* and the temperature-sensitive GAL4 repressor *tub-Gal80*[*ts*], hereafter referred to as *esgTS* (Micchelli and Perrimon, 2005). RNAi expression was induced by shifting the resulting young adults to 29°C, and intestinal tissue was analyzed seven days later. From this analysis, 21 genes led to aberrant nuclear poly(A)+ RNA accumulation and were grouped into two categories: strong accumulation (7 genes) and mild accumulation (14 genes). Furthermore,19 genes showed mild to severe progenitor cell loss and 16 of those also showed nuclear poly(A)+ RNA accumulation. Among these 16 genes, we focused on genes that showed both strong nuclear poly(A)+ accumulation and severe progenitor cell loss. Among these, the knockdown of *second mitotic wave missing* (*swm*) in progenitor cells led to a large accumulation of nuclear poly(A)+ RNA in progenitor cells (fig. 1C-D’). Swm is an understudied RBP that is predicted to have four known sequence motifs; a CCCH-type Zn^+^ finger domain, an RNA binding RNA recognition motif (RRM), and two nuclear localization sequences (Casso et al., 2008). It is implicated in nuclear RNA adenylation and export (Hurt et al., 2009; Yamamoto-Hino et al., 2010), and consistent with our findings in progenitor cells, its knockdown in S2R+ cells also leads to nuclear poly(A)+ RNA accumulation (Farny et al., 2008).

### Swm is expressed in nuclei of all cell types of the intestine

To determine where Swm was expressed in the adult intestinal epithelium, we generated a rat polyclonal antibody against the N-terminal 220 amino acids of Swm. The specificity of the resulting antisera was first validated by both western blotting (fig. S1B) and tissue immunostaining of *swm* mutant clones generated using the previously reported *swm^F14^* null allele (Casso et al., 2008) (fig. S1C and D). We then stained dissected wildtype adult female intestines with this antibody and found that Swm protein was expressed in all four cell types (ISCs, EBs, ees and ECs) (fig. 1E and fig. S1E). To carefully assess the sub-cellular localization of Swm in progenitor cells, we analyzed Swm expression in progenitor cells counterstained for cell membranes (myristoylated GFP), nuclear membranes (anti-Lamin antibodies), and nucleoli (anti-Fibrillarin antibodies). Swm was expressed exclusively in nuclei with no apparent staining in nucleoli (fig. 1F-G’’’’). This nuclear localization of the protein together with the accumulation of nuclear poly(A)+ RNA after *swm* knockdown, suggested a nuclear function for Swm.

### Loss of *swm* leads to progenitor cell loss

To assess the effects of *swm* loss in progenitor cells, we further analyzed the UAS-RNAi line identified in our screen (line HM05034 from the Transgenic RNAi Project (Ni et al., 2009)), which we refer to as *swm^RNAi-1^* for the remainder of this study. We crossed it to *esgTS* and analyzed intestines for progenitor cell number as well as total cell number after 1, 2, 5 and 10 days of RNAi. Progenitor cell number in *swm*-depleted intestines gradually declined over time and was nearly absent at the10-day timepoint (fig. 2A-C). Total cell number also declined with time (fig. S2A). We confirmed the loss of Swm in the *swm* RNAi-1 strain by staining for Swm (fig. S2B-C’) and verified the Swm mediated progenitor cell loss phenotype using a second RNAi strain, called *swm*^RNAi-2^, that we generated (fig. S2D).

**Figure 2.**
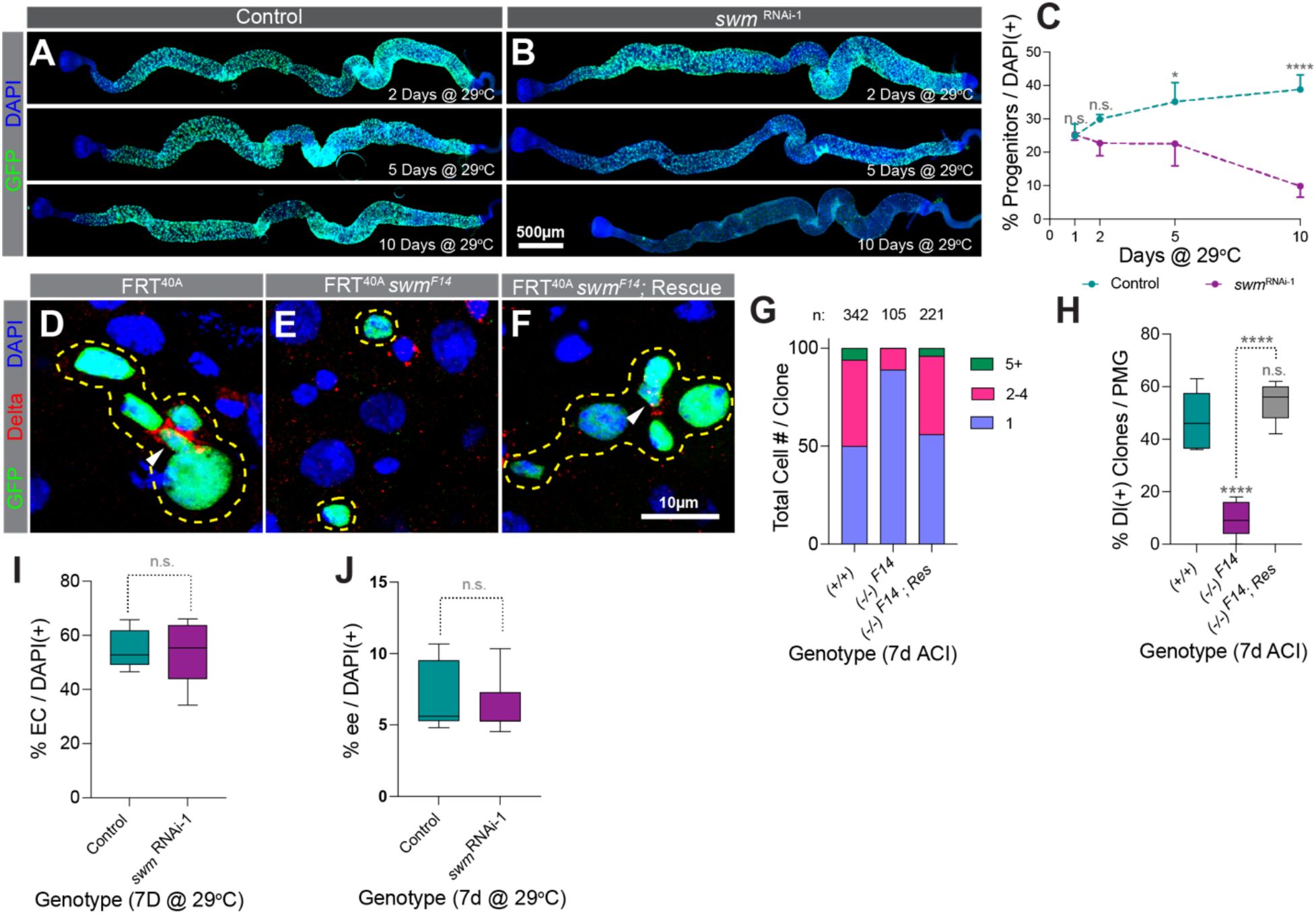
Depletion of *swm* results in loss of intestinal progenitor cells. (A and B) *esg*^TS^ (A) and *esg*^TS^; *swm* ^RNAi-1^ (B) intestines after 2, 5 and 10 days at 29°C. Ps are labelled by GFP (anti-GFP in green) and nuclei by DAPI (blue). (C) Normalized progenitor percentage (n=7 or 8) of genotypes shown in A and B after 1, 2, 5 and 10 days at 29°C. (D-F) Representative images of *tub-GAL4*, *UAS-GFP*-labeled (D) control, (E) *swm^F14^* and (F) rescued *swm^F14^* homozygous clones stained for Delta (red), GFP (green) and DAPI (blue). Clones are outlined in yellow and ISCs are indicated with white arrow heads. (G) Binned bar plot showing the quantification of total cell number per clone and (H) box and whisker graph showing percentage of Dl+ ISC containing clones per PMG in *tub-GAL4*, *UAS-GFP*-labeled control, *swm^F14^* and rescued *swm^F14^* intestines analyzed 7 days ACI. n indicates the number of clones analyzed for each genotype. (I and J) Box and whisker graphs showing (I) normalized EC percentage (n=7 or 8) in *mex1-GAL4*^TS^ and *mex1-GAL4*^TS^; *swm* ^RNAi-1^ and (J) normalized ee percentage (n=7 or 8) in *prosV1-GAL4*^TS^ and *prosV1-GAL4*^TS^ / *swm* ^RNAi-1^ intestines analyzed after 7 days at 29°C. Error bars on plots show mean±s.d. and asterisks denote statistical significance from Kruskal-Wallis test (C), ordinary one-way ANOVA (H) and Unpaired t-test (I and J). * p<0.05; ** p<0.01; *** p<0.001; **** p<0.0001; n.s., not significant. Complete genotypes are listed in table S1. ACI, after clone induction; EC, enterocyte; ee, enteroendocrine cell; Dl, delta; PMG, posterior midgut.

To further validate these results, we generated *swm ^F14^* mutant cell clones with the Mosaic Analysis with a Repressible Cell Marker (MARCM) technique (Lee and Luo, 1999) and analyzed intestines seven days after clone induction. Consistent with the *swm^RNAi-1^* RNAi results, 89% of *swm^F14^* mutant cell clones were single cell clones and the rest of the clones contained 2 or 3 cells (11%), while 50% of control clones contained single cells, 44% contained 2 to 5 cells, and 6% contained 6 or more cells. In addition, while 47% of control clones contained at least one cell stained for the ISC marker Dl (Ohlstein and Spradling, 2007), only 10% of *swm^F14^* mutant contained an ISC (fig. 2D, E, G and H). These defects were completely rescued by a transgenic insertion of a GFP-tagged ∼48kb fosmid clone that contained the entire *swm* locus (*fTRG01287.sfGFP-TVPTBF*) (fig. 2F-H) (Sarov et al., 2016). In contrast to depletion of *swm* in progenitor cells, knocking down *swm* in either ECs using the EC driver *midgut expression1-GAL4*, *mex1-GAL4* (Phillips and Thomas, 2006) or ee cells using *prosV1-GAL4,* an enhancer trap generated by the insertion of P{GawB} upstream of the transcription start site of *pros* (Balakireva et al., 1998), did not affect the respective differentiated cell number (fig. 2I, J, S2E-H’). Taken together, our data suggested that *swm* is specifically required in intestinal progenitor cells for their maintenance and normal tissue homeostasis.

### Loss of *swm* leads to loss of ISC/EB activity and identity

To determine whether ISCs can proliferate in the absence of *swm*, we induced ISC proliferation by feeding flies bleomycin, a DNA-damaging agent known to trigger ISC division (Amcheslavsky et al., 2009). *EsgTS*/*swm^RNAi-1^* flies were reared at the non-permissive temperature (18°C) until 4 days after eclosion, at which point flies were shifted to the permissive temperature (29°C) for an additional 2, 3, or 5 days. Flies were fed with bleomycin for the 24 hours prior to dissection. The number of mitotically active ISCs were then counted after immunostaining for phosphorylated-Histone 3 (pH3), a highly specific marker of condensed chromosomes during mitosis. Consistent with the *swm* mutant cell phenotype of single cell clones with very low-to-no Dl+ cells, knockdown of *swm* caused a significant reduction in the number of pH3 positive cells over time (fig. 3A). This could be due to the loss of the ability of ISCs to divide or the loss of ISCs themselves. To distinguish between these possibilities, we examined the cell identities of GFP+-progenitor cells expressing *swm^RNAi-1^* for 1, 3 and 5 days that were stained for both the Dl antibody, which labeled ISCs, as well as two previously described transgenic reporters that labeled EBs with *gbe-smGFP.V5.nls* and all progenitors with *mira-His2A.mCherry.HA* (Buddika et al., 2021a). Interestingly, we found that after 5 days of *swm^RNAi-1^* expression, nearly 80% of GFP+ cells did not express either the ISC-marker, Dl, or the EB-marker, *gbe-smGFP.V5.nls*, but did express the progenitor marker, *mira-His2A.mCherry.HA*, suggesting at least partial loss of ISC and EB cell identities (fig. 3B-D).

**Figure 3.**
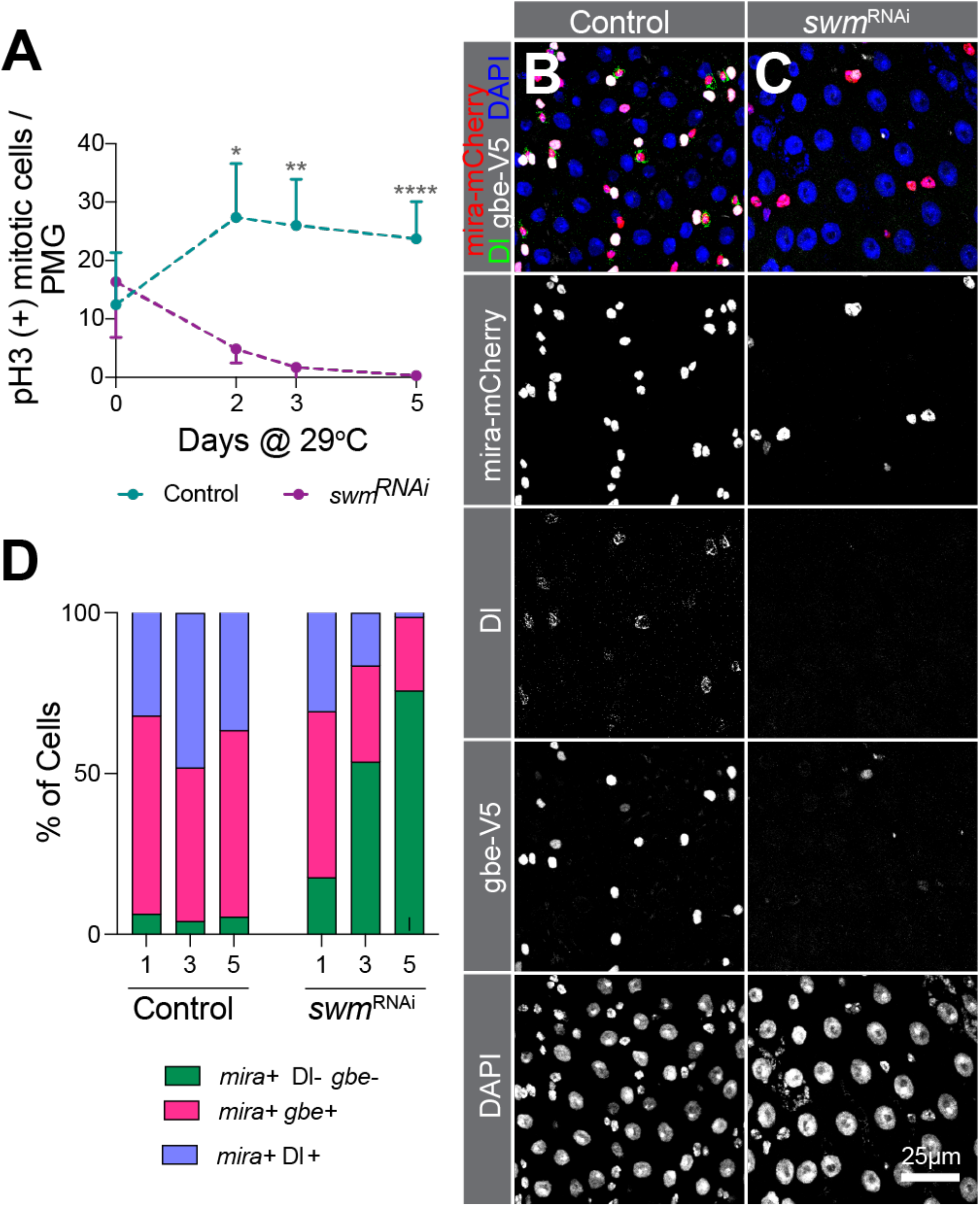
Loss of *swm* leads to loss of ISC/EB activity and identity. (A) Normalized pH3 (+) cell number per PMG (n=7 or 8) of bleomycin fed *esg*^TS^ and *esg*^TS^; *swm* ^RNAi-1^ intestines after 0, 2, 3 and 5 days at 29°C. (B-D) PMG section from control (B) and *swm* ^RNAi-1^ (C) after 5 days at 29°C stained for Ps (anti-RFP for mira-mCherry, in red), ISCs (anti-Dl, in green), EBs (anti-V5 for gbe-smGFP.V5.nls, in white) and all nuclei (DAPI, in blue). Individual channels are shown in gray scale. (D) Binned bar plot showing the quantification of percentage of mira+Dl-gbe-(no defined identity), mira+gbe+ (EB), and mira+Dl+ (ISC) of intestines from genotypes shown in B and C after 1, 3 and 5 days at 29°C. Error bars on plots show mean±s.d. and asterisks denote statistical significance from Kruskal-Wallis test (A). * p<0.05; ** p<0.01; *** p<0.001; **** p<0.0001; n.s., not significant. Complete genotypes are listed in table S1. P, progenitor cell; ISC, intestinal stem cell; EB, enteroblast; Dl, delta; PMG, posterior midgut.

One possible reason for the progenitor cell number reduction could be due to accelerated differentiation of ISCs into ECs or ee cells. To test whether *swm* depleted progenitors differentiate into ECs and ee cells, we took advantage of the Repressible Dual Differential stability cell Marker (ReDDM) cell-lineage tracing system (Antonello et al., 2015). This system uses two reporters, a short-lived GFP and a longer-lived, highly stable histone tagged RFP, both of which are under the regulation of the *esg*-GAL4 driver. Undifferentiated progenitors are labelled with both GFP and RFP but, since GFP degrades as cells differentiates, the newly differentiated ECs and ee cells are only labelled with RFP. ReDDM analysis after 10 days of *swm* RNAi revealed that, in the absence of *swm*, the percentage of newborn ECs and ee cells was largely reduced (4%) compared to the control (41%) (fig. S3A-E), suggesting that *swm* depleted progenitor are severely compromised in their differentiation into EC and ee cells.

Moreover, this result indicates that the severe loss of stem cells in *swm* mutants is not because they inappropriately differentiate into EC and ee. Taken together, these data indicated that *swm* is required for ISC and EB cell identity and thereby homeostasis of differentiated cell numbers.

### *swm* depleted progenitor cells detach from the basement membrane

Why are progenitor cells lost in the absence of *swm*? One possible cause of cell loss is cell death. To test whether *swm* depleted progenitor cells undergo apoptotic cell death, we used ApopTag assay and found that progenitor cells were not ApopTag positive (fig. S3F). Furthermore, the cell loss phenotype was not rescued by overexpression of the apoptosis inhibitor P35, further suggesting that apoptosis was not the cause of cell loss (fig. S3G). During this analysis, we noticed that *swm* depleted progenitor cells appeared small and round in comparison to the control progenitor cells, which were mostly triangular (fig. 4A and B). One possible reason for the altered cell shape is due to disruption of the physical attachment of progenitor cells to the underlying basement membrane; this basal location of progenitor cells is likely important for proper reception of signals for their maintenance and function (Lin et al., 2013). To determine whether the progenitor cell-basement membrane attachment was affected, we took a confocal microscopy-based approach where we labeled the Filamentous-actin (F-actin) of intestinal tissue with phalloidin and progenitors with GFP, collected images of Z sections spaced 0.3µm from one another that spanned the basal-to-lumen sides of the intestine, and reconstructed three-dimensional models from them. After five days of *swm^RNAi-1^* expression, within a total population of 94 cells, we observed three different GFP+ progenitor cell populations: 1) one that was fully detached from the basement membrane and shifted towards the lumen side (42% of cells), 2) one that was partially detached from the basement membrane (28% of cells), and 3) one that was fully attached (30% of cells). In comparison, 100% of control progenitor cells (n=54) were fully attached to the basement membrane (fig. 4C-F, supplementary video 1-3).

**Figure 4.**
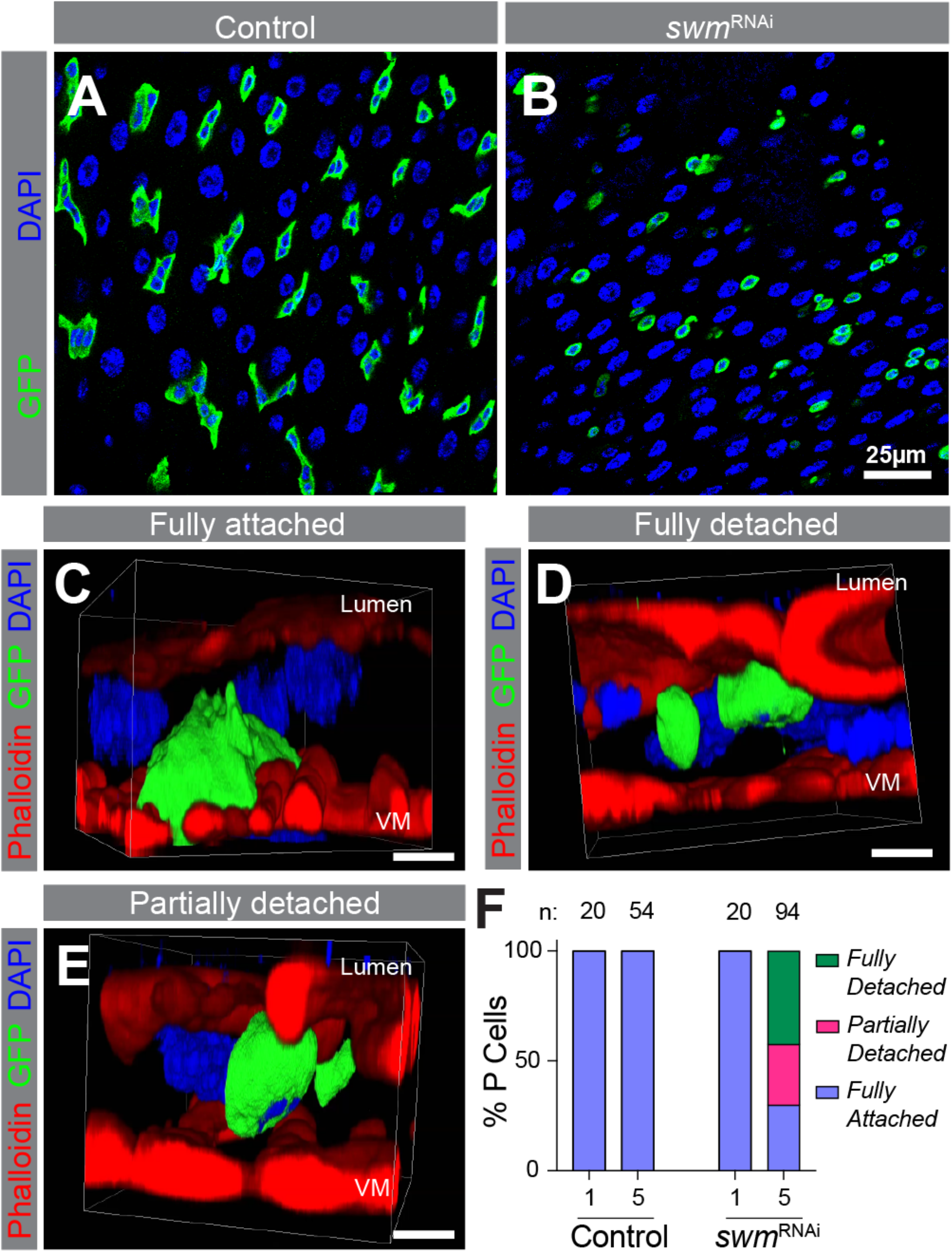
In the absence of swm, Intestinal progenitors detach from the basement membrane. (A and B) PMG section from *esg*^TS^ (A) and *esg*^TS^; *swm* ^RNAi-1^ (B) after 5 days at 29°C. Ps are labelled by GFP (anti-GFP in green) and nuclei with DAPI (blue). (C-E) Representative snapshot of 3-D reconstructed confocal images of fully attached cell from *esg*^TS^ (C), fully detached cell from *esg*^TS^; *swm* ^RNAi-1^ and partially detached cell from *esg*^TS^; *swm* ^RNAi-1^ (E) after 5 days at 29°C labelled F-actin (phalloidin-TRITC in red), Ps (anti-GFP in green) and nuclei (DAPI in blue). Apical lumen side is on top and basal VM is on the bottom of images as labelled. (F) Binned bar plot showing the quantification of fully detached, fully attached, and partially detached P cell percentages in *esg*^TS^ control (left) and *esg*^TS^; *swm* ^RNAi-1^ (right) after 1 and 5 days at 29°C. n indicates the number of individual cells used for the quantification across up to 5 intestines in each time point. Complete genotypes are listed in table S1. VM, visceral muscle; P, progenitor cell.

Taken together, our data indicated that in the absence of *swm*, progenitors detached from the basement membrane and shifted towards the lumenal side of the gut tissue, likely leading to progenitor cell elimination from the epithelium.

### Swm physically associates with epithelial cell maintenance and adhesion related transcripts

To begin to understand the mechanism of Swm function, we set out to identify the mRNAs that are bound by Swm using crosslinking immunoprecipitation (CLIP) combined with sequencing (CLIP-seq). Using the anti-Swm antibody, endogenous Swm was immunoprecipitated from lysates prepared from UV-crosslinked, dissected intestines of *w^1118^* females (fig. S4A). We prepared libraries from the RNA extracted from two independent immunoprecipitates (Swm-IPs) and, in parallel, a third library was prepared from RNA extracted from one ‘input’ lysate, which we used as our normalization control. High-throughput sequencing and bioinformatic analysis identified 418 genes enriched in the Swm-IP samples compared to the input lysate (fig. 5A and Table S3). We validated three Swm-IP enriched targets (*cheerio* [*cher*]*, headcase* [*hdc*], and *CG12194*) by performing qPCR on Swm-IPs, thus demonstrating that our results were reliable (fig. S4B). Interestingly, Gene Ontology (GO) analysis showed that epithelial cell differentiation, actin cytoskeleton organization, cell fate commitment, and cell adhesion related genes were enriched in Swm-IPs (fig. 5B). Genes such as *cher*, *coracle* (*cora*), *discs large 1* (*dlg1*), *futsch*, *molecule interacting with CasL* (*mical*), *Myosin 10A* (*Myo10A*), *rhea*, *scribble* (*scrib*), *short stop* (*shot*), *split ends* (*spen*), *Tenascin accessory* (*Ten-a*), and *Tenascin major* (*Ten-m*) were found among SWM-IP enriched genes (Table S3). Taken together, our identification of RNAs associated with endogenous Swm in intestinal tissue suggested a role for Swm in regulating genes involved in epithelial cell adhesion, cell fate commitment and maintenance.

**Figure 5.**
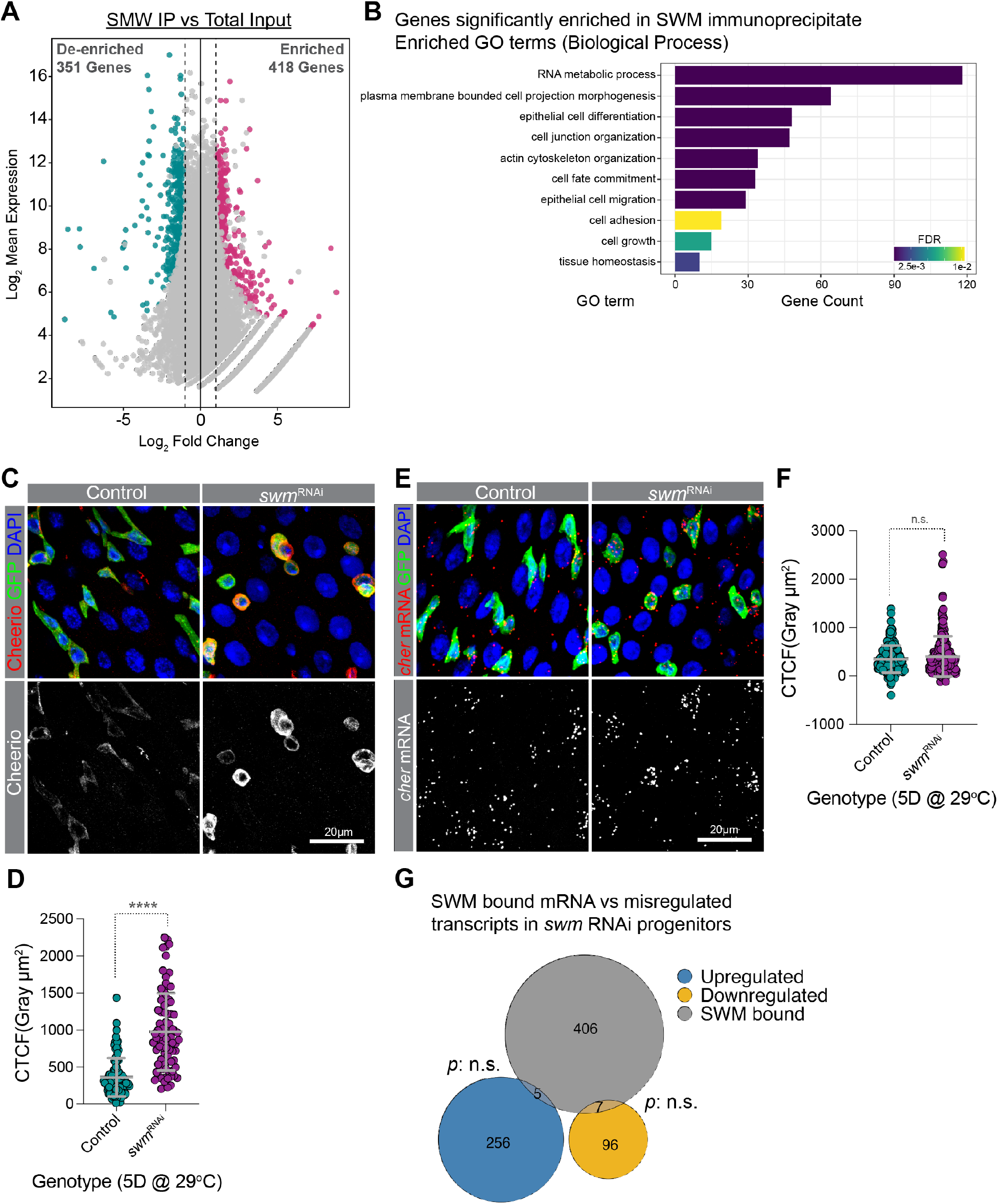
Swm binds and post-transcriptionally regulates transcripts related to cell adhesion. (A) A scatter dot plot visualizing differentially enriched genes in SWM IP versus total input. Each dot represents a single gene. Cyan and pink dots indicate genes with a false discovery rate (FDR) adjusted p-value < 0.05 and a Log_2_ fold change < −1 or > 1, respectively. (B) A bar plot showing a selected set of significantly enriched Gene Ontology (GO) terms (biological processes) for genes that are physically associated with Swm. (C) Representative confocal micrographs from PMGs of *esg*^TS^ control (left) and *esg*^TS^; *swm* ^RNAi-1^ (right) after 5 days at 29°C stained for Cheerio (red) and DAPI (blue). Ps are labelled with GFP in green. Bottom panel shows Cheerio staining in gray scale. (D) Scatter dot plot of normalized Cheerio fluorescence intensity (corrected total cell fluorescence, CTCF) of progenitor cells from *esg*^TS^ (n=95 cells, 8 intestines) and *esg*^TS^; *swm* ^RNAi-1^ (n=84 cells, 7 intestines) after 5 days at 29°C. (E) Intestinal progenitors from PMGs of *esg*^TS^ control (left) and *esg*^TS^; *swm* ^RNAi-1^ (right) after 5 days at 29°C labelled cheerio mRNA (red), Ps stained for GFP (anti-GFP in green) and nuclei with DAPI (blue). Bottom panel shows cheerio mRNA in gray scale. (F) Scatter dot plot of normalized cheerio mRNA fluorescence intensity (CTCF) of progenitor cells from genotypes in F (n=167 cells, 7 intestines for *esg*^TS^ and n=241 cells, 10 intestines for *esg*^TS^; *swm* ^RNAi-1^). (G) Venn diagram showing the overlap between genes enriched in Swm-CLIP (gray) and genes that are upregulated (blue) or downregulated (yellow) in *swm* depleted progenitor cells identified from RNAseq. Error bars on plots show mean±s.d. and asterisks denote statistical significance from Mann-Whitney test (D, F). * p<0.05; ** p<0.01; *** p<0.001; **** p<0.0001; n.s., not significant. Complete genotypes are listed in table S1. PMG, posterior midgut; P, progenitor cell; CTCF, corrected total cell fluorescence.

### Swm post-transcriptionally regulates target mRNAs

To begin to understand how Swm binding to transcripts impacted expression of genes involved in epithelial differentiation, actin cytoskeleton organization, cell fate commitment, and cell adhesion, we knocked down *swm* in progenitor cells using RNAi for five days and immunostained for proteins encoded by Swm-bound transcripts using available antibodies. These included a total of nine proteins encoded by *Cadherin-N* (*CadN*), *cher*, *cora*, *cut* (*ct*), *dlg1*, *hdc*, *osa*, *rhea*, and *shot*. Among the proteins for which we stained, Cheerio (encoded by *cher*), Talin (encoded by *rhea*) and Shortstop (encoded by *shot*) were noticeably upregulated, and Cheerio showed the most obvious increase in progenitor cells upon *swm* loss (fig. 5C and D). Cheerio is a *Drosophila* member of the Filamin family of actin-binding proteins (Sokol and Cooley, 1999).

Filamins bind to F-actin to crosslink actin filaments into parallel bundles and connect F-actin to cell membranes, and have been shown to play key roles in modulating cell shape, cell adhesion, cell motility, and differentiation (Lamsoul et al., 2020; Sokol and Cooley, 1999). In addition to Cheerio, Talin protein was also increased in *swm* depleted progenitor cells (fig. S4C-E). Talin is a large adaptor protein, encoded by the gene *rhea,* which links ECM-bound integrins to the actin cytoskeleton and thereby regulates adhesion of cells to the ECM (Brown et al., 2002).

Given the upregulated protein levels, to begin to understand how Swm was mediating effects on these bound RNAs, we next asked whether transcript level or distribution were affected. To address this question, we analyzed the subcellular localization of *cher* transcripts in *swm* depleted intestinal progenitors using RNAscope *in situ* hybridization probes after first verifying the specificity of *cher* probes in progenitors expressing *cher* RNAi (fig. S5A-D’) (Wang et al., 2012). From the analysis, we found that *cher* transcript level was not significantly changed in GFP+ cells after five days of *swm* RNAi in progenitor cells (fig. 5E and F) even though our previous analysis indicated that Cher protein was elevated at this timepoint. In addition, we did not observe an obvious change in cellular distribution of the transcript. To investigate the steady state mRNA levels of Swm-IP targets after *swm* depletion, we performed transcriptomic profiling (RNA-seq) of FACS (fluorescent activated cell sorting) isolated *swm* RNAi and control intestinal progenitor cells after five days of RNAi. In agreement with the RNAscope *in situ* hybridization results, *swm* knock down in progenitor cells did not significantly alter the abundance of Swm-IP target transcripts, including *cher*, and, despite the loss of progenitor cell identity and functions we had observed, the transcriptome of *swm^RNAi-1^*-expressing progenitor cells showed only slight changes compared to control cells (fig. 5G, Table S4). Altogether, the analysis of the Cher protein and transcript levels in *swm*-depleted progenitor cells indicated that Swm post-transcriptionally regulated the levels of Cher and likely other proteins encoded by Swm-bound transcripts.

## DISCUSSION

Here we report the results of a candidate RNAi screen that identified the RBP Swm as critically important to maintain ISC and EB cell identities and thereby progenitor cell function. This conclusion is based on observations that Swm is nuclear localized; that nuclear poly(A)+ RNA accumulates in the absence of Swm; that Swm depleted progenitor cells lose physical contact with the basement membrane, defined cell identities and are lost from the epithelium; that Swm physically associates with mRNAs related to epithelial cell maintenance and adhesion including cher; and, finally, that Swm loss leads to upregulation of proteins encoded by Swm-bound targets, including Cher, without affecting transcript levels.

Based on our results, we propose a model where Swm binds and regulates the translation of mRNAs involved in maintaining epithelial progenitor cell identity and function. An open question is how exactly Swm regulates the expression of its bound RNA targets. We speculate that Swm regulates the poly(A) tail length of the mRNAs, that these mRNAs are hyperadenylated in the absence of Swm, and that hyperadenylation of targeted mRNAs leads to enhanced protein translation in intestinal cells. Supporting this, longer poly(A) tails can improve protein translation in a context dependent manner in both *Drosophila* and vertebrate models (Eichhorn et al., 2016; Passmore and Tang, 2021; Xiang and Bartel, 2021). In addition, Swm has been reported to be a physical interactor of *Drosophila* Zinc finger CCCH domain-containing protein 3 (dZC3H3), and depletion of dZC3H3 in *Drosophila* S2R+ cells resulted in hyperadenylation of mRNAs (Hurt et al., 2009). Importantly, dZC3H3 also interacts with Polyadenylate-Binding Protein 2 (Pabp2), the *Drosophila* nuclear poly(A) binding protein, which controls poly(A) tail length. Our RNAi screen revealed that progenitor specific knock down of *dZC3H3*, just like *swm*, resulted in severe nuclear poly(A)+ RNA accumulation and cell loss (Table S1), supporting the notion that these genes also functionally interact in intestinal tissue. The two human orthologs of Swm, RNA Binding Motif protein 26 and 27 (RBM26 and RBM27), are also known to bind to ZC3H3 in HEK293 cells (Silla et al., 2020), indicating that the functional interaction between these proteins is evolutionarily conserved. Furthermore, despite the nuclear poly(A)+ RNA accumulation in *swm* depleted progenitor cells, we did not observe a nuclear accumulation of *cher* mRNA by *in situ* hybridization. One possible reason for this could be that one RNA target is not enough to visualize the accumulation, but rather the accumulation is a result of multiple poly(A)+ RNAs. Another explanation is that the increase in nuclear poly(A)+ signal can be a result of the hyperadenylation of transcripts but not an increase in their numbers of copies, explaining why it was detected by oligo d(T) but not *cher* mRNA *in situ* probes.

One of the Swm mRNA targets that we identified encodes Cher, a member of the Filamin family of actin binding proteins. Mammalian Filamin A is known to control the architecture and mechanics of the actin cytoskeleton in a dosage-dependent manner. Tighter F-actin bundles were observed at high Filamin A concentration, leading to reduced flexibility of cells. In contrast, at lower Filamin-A levels, the F-actin cytoskeleton was more dynamic. Filamin A is also known to negatively regulate cell adhesion by binding to the cytoplasmic domains of integrins (Lamsoul et al., 2020). It is possible that modulation of Cher levels could adjust progenitor cell adhesion properties that contribute to the maintenance of stem cell identity. Given the large number of Swm-bound mRNAs that we identified, however, we suspect that Swm likely regulates multiple mRNA targets coordinately to adjust stem cell adhesion. Consistent with this idea, RNAi knockdown of Cher did not rescue the *swm* loss-of-function phenotype (data not shown), suggesting that elevated Cher levels are not the only cause of *swm* phenotypes.

Even though Swm is expressed in all four cell types of the intestinal epithelium, only progenitor cells were lost in the absence of *swm*. One likely reason for this differential requirement of Swm for cell survival could be the differential requirement of Swm-bound target mRNAs for the viability of the different cell types of the intestine.

Among Swm-IP targets, genes encoding cell adhesion-related molecules, including integrin linker molecules and actin cytoskeleton binding molecules, were abundant. Adult stem cells are typically associated with a specific microenvironment or niche and this stem cell attachment is frequently mediated via integrins, integrin binding linker molecules, and the cellular actin cytoskeleton (Chen et al., 2013; Xi, 2009). Recent single cell transcriptomic analyses, tissue immunostaining, and mutant studies have shown that cell adhesion-related molecules are differentially expressed and required for intestinal progenitors, including both integrins, encoded by *myospheroid* (*mys*), *multiple edematous wings* (*mew*) and scab (*scb*), as well as integrin linker molecules, such as *talin*/*rhea*, *Laminin A* (*lanA*), and *integrin-linked kinase* (*ilk*) (Hung et al., 2020; Lin et al., 2013).

We found that *swm* depleted progenitor cells do not express Dl or *gbe*, indicating a loss of ISC and EB cell identities. Identities in the *Drosophila* intestinal cell lineage are specified by Notch signaling (Micchelli and Perrimon, 2005; Ohlstein and Spradling, 2005; Ohlstein and Spradling, 2007). ISCs express the Notch ligand, Dl and thereby activate Notch signaling in neighboring ISC daughter cells, EBs. Downregulation of Dl in *swm*-depleted ISCs disrupts Notch signaling and that might compromise EB cell identity and subsequent cell differentiation. However, whether this *swm*-mediated Dl downregulation is the cause, or the effect of progenitor cell detachment is an open question. Furthermore, although other studies have shown that loss of Dl leads to loss of Notch signaling and induces ee-like tumors (Ohlstein and Spradling, 2007), *swm* loss did not result in a tumor-like cell mass. This could be simply due to loss of ISC identity including impaired cell proliferation when *swm* is absent. A similar phenotype was reported for *terribly reduced optic lobes* mutant clones, which lost Dl but did not display an overproliferation phenotype (You et al., 2014).

Another open question raised by our study is how *swm* mutant progenitor cells are lost from the epithelium. Previous studies have found that stem cells that detach from the ECM undergo anoikis, a type of apoptosis that is commonly triggered by loss of cell adhesion (Gilmore, 2005). However, we found that detached progenitors do not undergo apoptosis, a finding consistent with other studies in the *Drosophila* intestines that showed that the progenitor cell loss that was associated with the loss of integrins, integrin linker molecules, or ECM components was not caused by apoptosis (Lin et al., 2013; You et al., 2014). These observations indicate that intestinal progenitors are not generally eliminated by anoikis. However, we cannot rule out the possibility that detached progenitor cells are eliminated via a different cell death mechanism. As an alternative, we hypothesize that detached or extruded progenitor cells are shed into the lumen and thereby get eliminated from the epithelium. Intestinal progenitor cell extrusion has not been previously reported, but live imaging of the midgut has revealed enterocyte extrusion from the epithelium (Martin et al., 2018), and it has also been reported that ISC tumors promote apoptotic enterocyte extrusion to create space on the basement membrane (Patel et al., 2015).

Nuclear post-transcriptional gene regulatory mechanisms in stem cell biology are largely understudied (Wang et al., 2013) and our study suggests active nuclear post-transcriptional mechanisms in intestinal cells. Future work including identifying molecular and genetic interactors of Swm will dissect the Swm-mediated nuclear post-transcriptional regulatory mechanisms.

## Data Availability Statement

Strains and plasmids are available upon request. RNA-seq data will be submitted to GEO and an accession number will be included in a revised version of the manuscript prior to publication. Table S1 lists the full genotypes used in each figure. Table S2 lists genes screened by RNAi. Table S3 lists genes enriched in Swm-IPs. Table S4 lists differentially expressed genes identified by RNA-seq. Table S5 lists the reagent used in the study, including antibodies, fly strains, and oligos. Supplementary videos show the 3D reconstructions from Figure 4.

## Acknowledgements

We thank Lynn Cooley, Jean-Francois Ferveur, Claire Thomas, the Bloomington *Drosophila* Stock Center (supported by grant NIHP4OOD018537), the Vienna *Drosophila* Resource Center (VDRC), the *Drosophila* Genome Resource Center (supported by grant NIH2P40OD010949), and the Developmental Studies Hybridoma Bank (created by the NICHD of the NIH) for reagents; the FlyBase for resources and information; the Light Microscopy Imaging Center (supported by grant NIH1S10OD024988-01) for access to the SP8 confocal; the Flow Cytometry Core Facility at Indiana University, Bloomington for access to the BD FACSAria™ II flow cytometer; and the National Institute of General Medical Sciences (Award R01GM124220 to NSS and transferred to BRC) for financial support.

---

**Figure S1.**
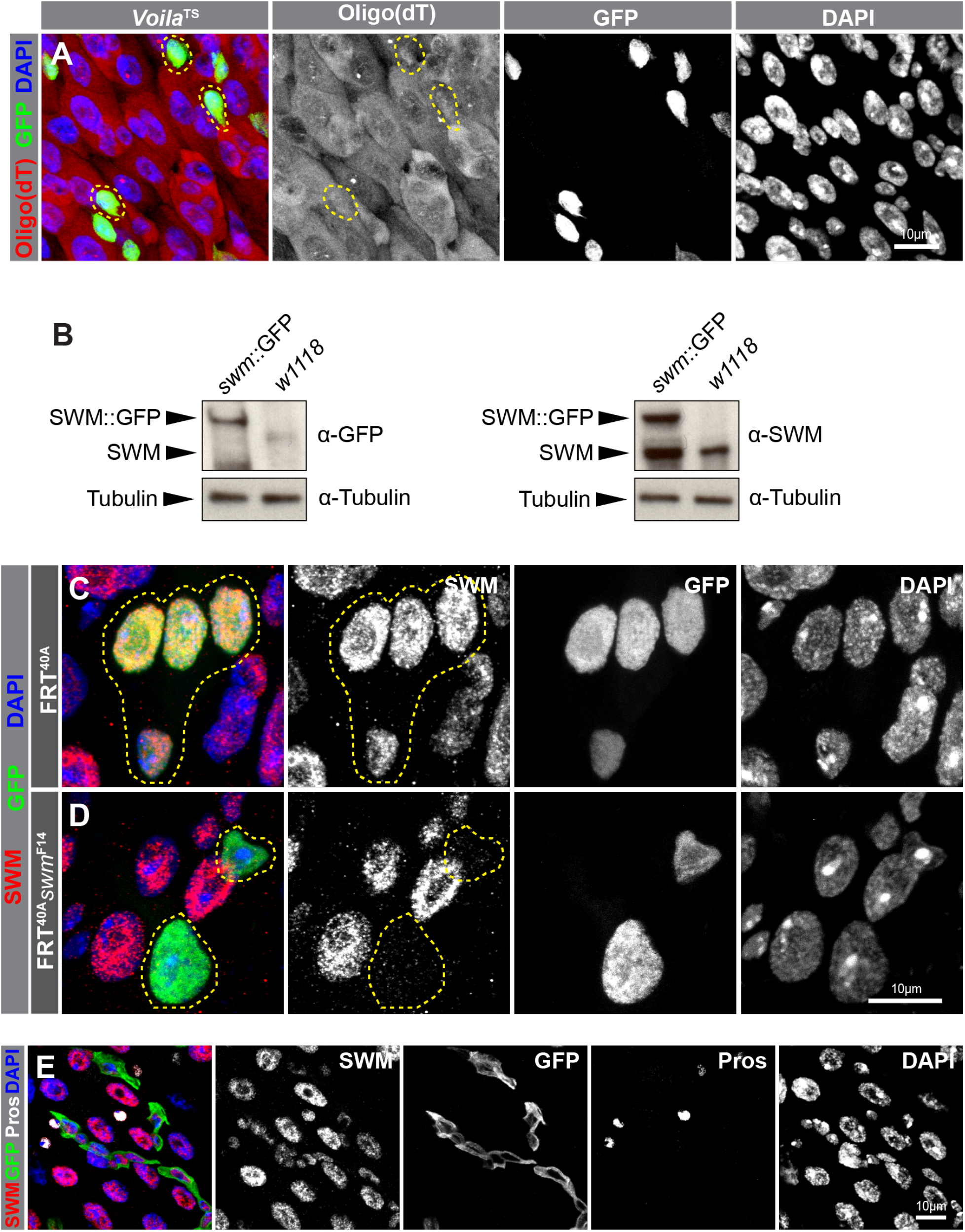
Poly(A)+ RNA distribution in ee cells and Swm antibody verification. (A) Poly(A)+ RNA distribution in cells from *prosV1-GAL4*^TS^ intestines, labelled by oligo (dT) probes (red) and ee cells are labelled by GFP (anti-GFP in green) and nuclei with DAPI (blue). Individual channels are shown in gray scale. (B-D) Validation of the specificity of the Swm antibody using western blotting and tissue immunostaining. (B) Western blots of w[1118] and swm-GFP/ + fly extracts probed with anti-GFP (left panel), anti-Swm (right panel) and anti-Tubulin (bottom panels) antibodies. (C-D) Representative images of *tub-GAL4*, *UAS-GFP*-labeled (D) control, (E) *swm^F14^* homozygous clones stained for Swm (anti-Swm, in red), GFP (anti-GFP, in green) and nuclei (DAPI, in blue). Control (C) and *swm* mutant (D) cells are outlined in yellow dashed line. Individual channels are shown in gray scale. (E) Individual channels of the PMG section shown in Figure 1D, stained for Swm (anti-Swm in red). Ps are shown in GFP (green), ee cells are stained for Prospero (anti-Prospero in white) and nuclei for DAPI (blue). Complete genotypes are listed in table S1. Ee, enteroendocrine cell; PMG, posterior midgut, P, progenitor cell.

**Figure S2.**
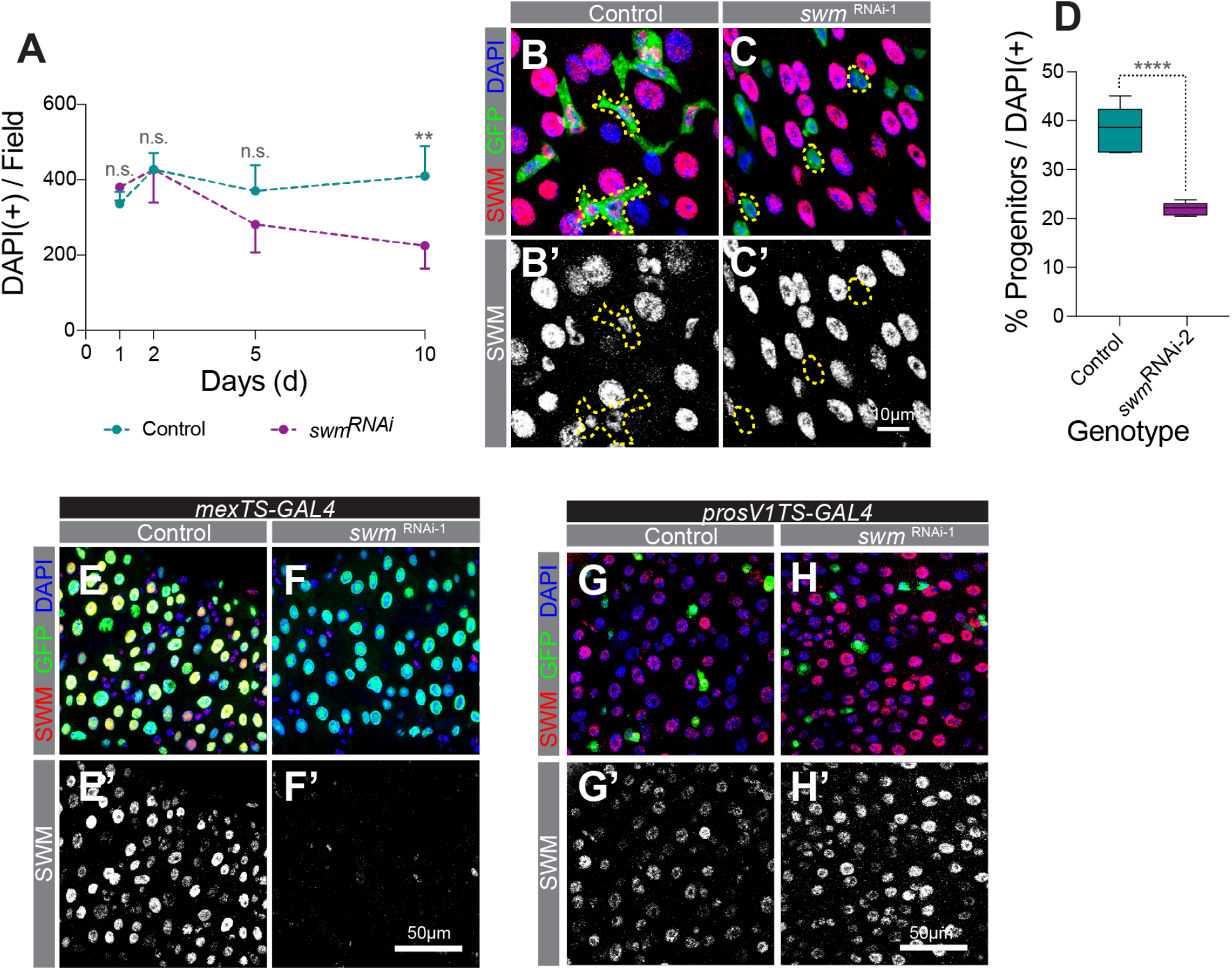
Loss of swm in ECs or ee cells does not affect respective differentiated cell number. (A) Normalized DAPI (+) cell number per field (n=7 or 8) of genotypes *esg*^TS^ and *esg*^TS^; *swm* ^RNAi-1^ shown in Figure 2A and 2B after 1, 2, 5 and 10 days at 29°C. (B-C) PMG sections from (B) *esg*^TS^ and (C) *esg*^TS^; *swm* ^RNAi-1^ after 5 days at 29°C, stained for Swm (anti-Swm, in red) and all nuclei (DAPI, in blue). Ps are shown in GFP (green). (D) Box and whisker graphs showing normalized P percentage (n=7 or 8) in *esg*^TS^ and *esg*^TS^; *swm* ^RNAi-2^ intestines analyzed after 7 days at 29°C. (E-F’) PMG sections from (E) *mex1-GAL*4^TS^ and (F) *mex1-GAL4*^TS^; *swm* ^RNAi-1^ intestines after 7 days at 29°C stained for Swm (anti-Swm, in red) and all nuclei (DAPI, in blue). ECs are shown in GFP-nls (green). Swm stained red channel of control (E’) and *swm* ^RNAi-1^ (F’) is shown in gray scale. (G-H’) PMG sections from (G) *prosV1-GAL4*^TS^ and (H) *prosV1-GAL4*^TS^; *swm* ^RNAi-1^ intestines after 7 days at 29°C stained for Swm (anti-Swm, in red) and all nuclei (DAPI, in blue). Ee cells are shown in GFP (green). Swm stained red channel of control (G’) and *swm* ^RNAi-1^ (H’) is shown in gray scale. Error bars on plots show mean±s.d. and asterisks denote statistical significance from Kruskal-Wallis test (A) and Unpaired t-test (D). * p<0.05; ** p<0.01; *** p<0.001; **** p<0.0001; n.s., not significant. Complete genotypes are listed in table S1. PMG, posterior midgut; P, progenitor cell; EC, enterocyte; ee, enteroendocrine cell.

**Figure S3.**
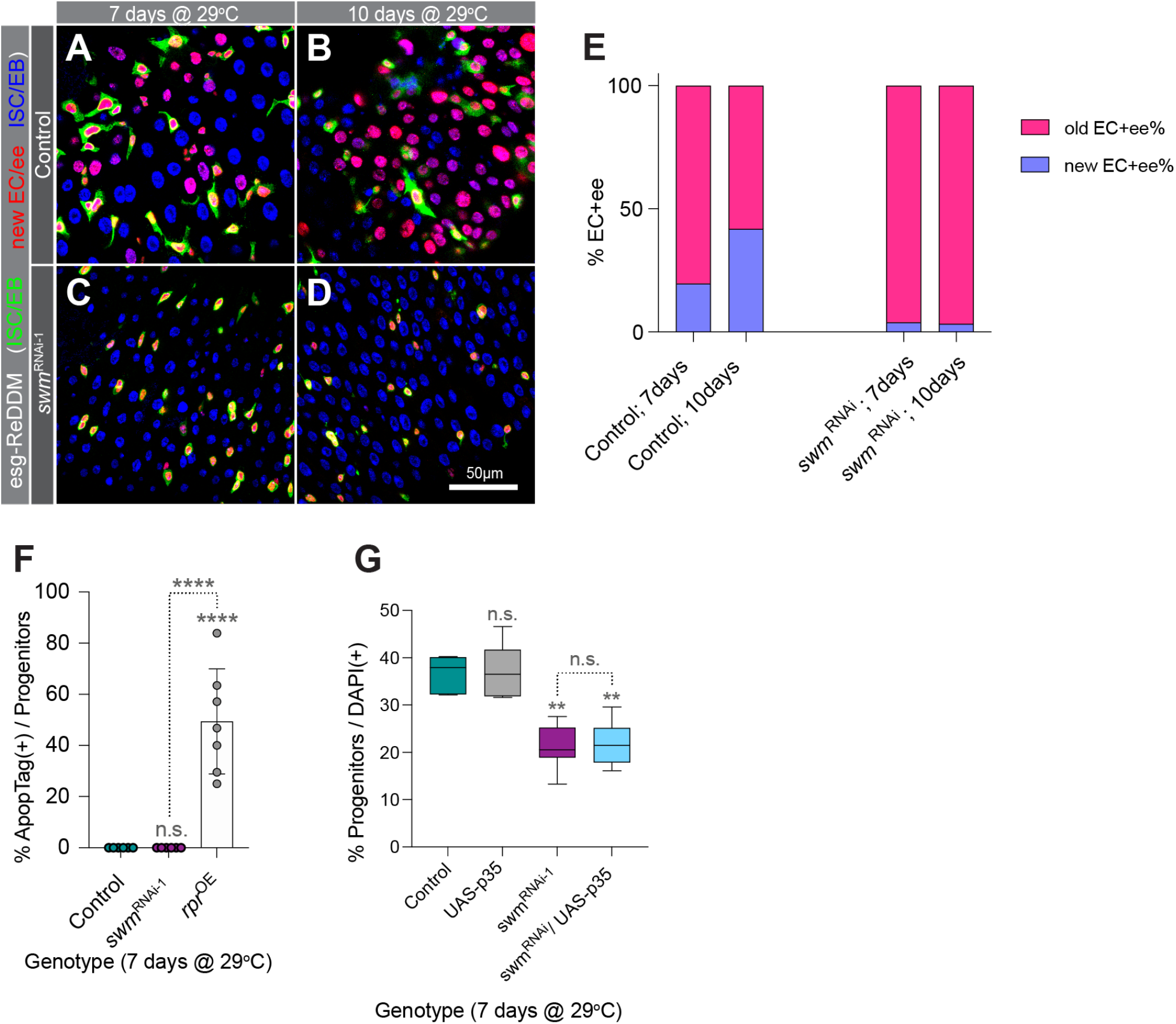
*swm* depleted progenitor cells do not produce differentiated cells and do not undergo apoptosis. (A-D) PMG sections of esg-ReDDM crossed to w[1118] (A and B) and to *swm* ^RNAi-1^ (C and D) intestines after 7 days at 29°C (A and C) and 10 days (B and D) stained for nuclei (DAPI, in blue). All Ps are labelled by both GFP and RFP (cytoplasmic green and nuclear red) and new ECs and ee cells are shown in only RFP (nuclear red). (E) Binned bar plot showing the quantification of percentage of old EC+ee and new EC+ee of intestines from genotypes shown in A-D panels (n=6 intestines). (F) Bar graph showing normalized percentage of ApopTag+ cells of intestines from control (*esg*^TS^), *esg*^TS^; *swm*^RNAi-1^ and *esg*^TS^; *rpr^OE^* after 7 days at 29°C (n=7 or 8 intestines). (G) Box and whisker graph showing normalized percentage of Ps of intestines from control (*esg*^TS^), *esg*^TS^; *UAS-p35*, *esg*^TS^; *swm*^RNAi-1^, and *esg*^TS^; *swm*^RNAi-1^/ *UAS-p35* after 7 days at 29°C (n=6 intestines). Error bars on plots show mean±s.d. and asterisks denote statistical significance from ordinary one-way ANOVA (F and G). * p<0.05; ** p<0.01; *** p<0.001; **** p<0.0001; n.s., not significant. Complete genotypes are listed in table S1. PMG, posterior midgut; P, progenitor cell; EC, enterocyte; ee, enteroendocrine cell; OE, overexpression; ReDDM, Repressible Dual Differential stability cell Marker.

**Figure S4.**
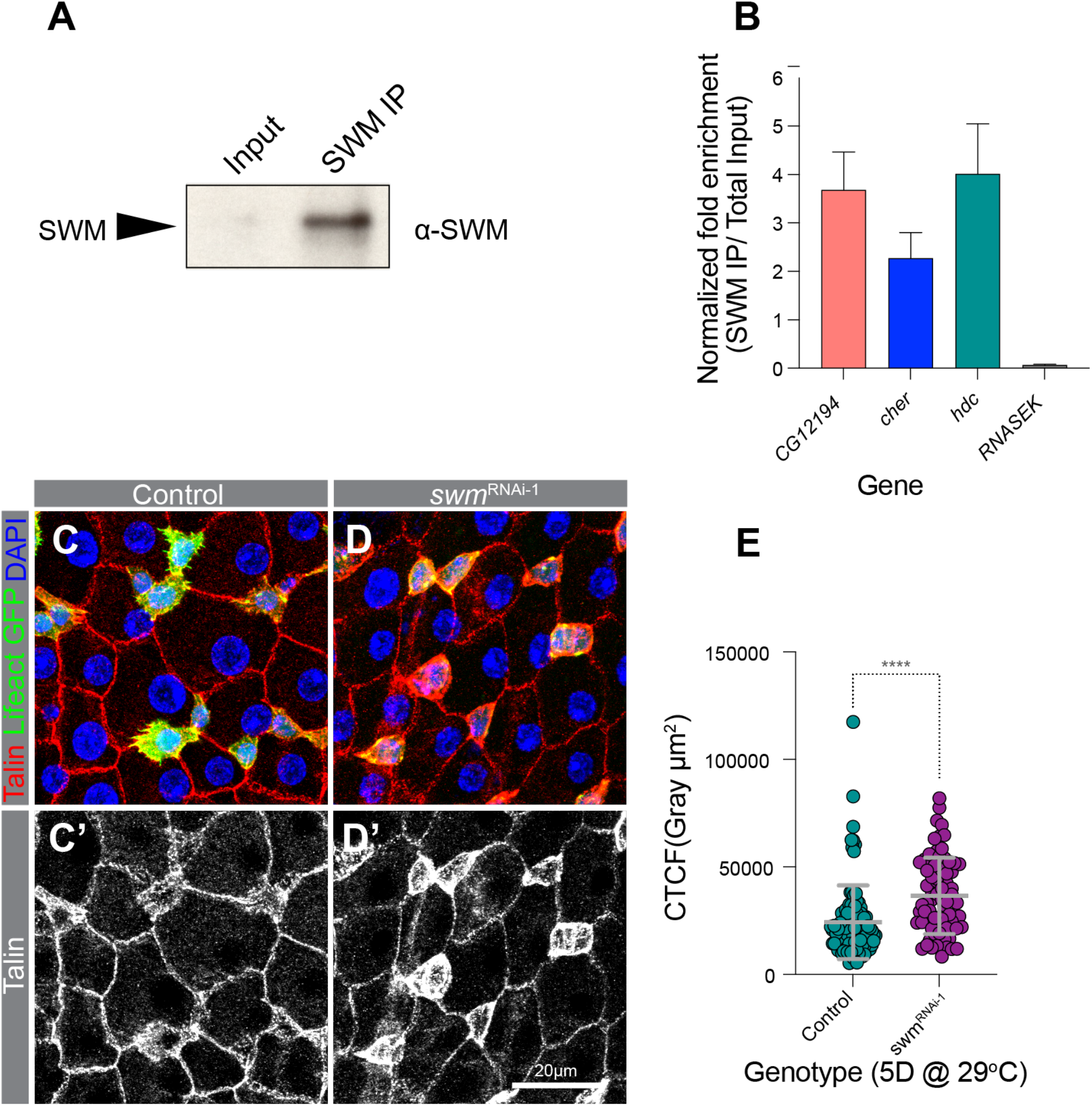
RIP-qPCR verification of Swm-IP targets and Talin expression after *swm* loss. (A) Western blot of Swm-IP and total input probed with anti-Swm antibody. (B) Normalized fold enrichment of *CG12194*, *cher*, *hdc*, and *RNASEK* transcripts in Swm-IP determined by qPCR. (C-D’) Sections of PMG from (C) control (*UAS-Lifeact-GFP*) and (D) *UAS-Lifeact-GFP*; *swm*^RNAi-1^ after 5 days at 29°C stained for Talin (anti-Talin, in red) and nuclei (DAPI, in blue). Ps are labelled with Lifeact-GFP (green). (C’ and D’) Anti-Talin staining is shown in gray scale. (E) Scatter dot plot of normalized Talin fluorescence intensity (corrected total cell fluorescence, CTCF) of progenitor cells from genotypes shown in C and D panels (n=97 cells, from 7 intestines for control and n=70 cells, from 8 intestines for *swm*^RNAi-1^). Error bars on plots show mean±s.d. and asterisks denote statistical significance from Mann-Whitney test (E). * p<0.05; ** p<0.01; *** p<0.001; **** p<0.0001; n.s., not significant. Complete genotypes are listed in table S1. PMG, posterior midgut; P, progenitor cell; CTCF, corrected total cell fluorescence.

**Figure S5.**
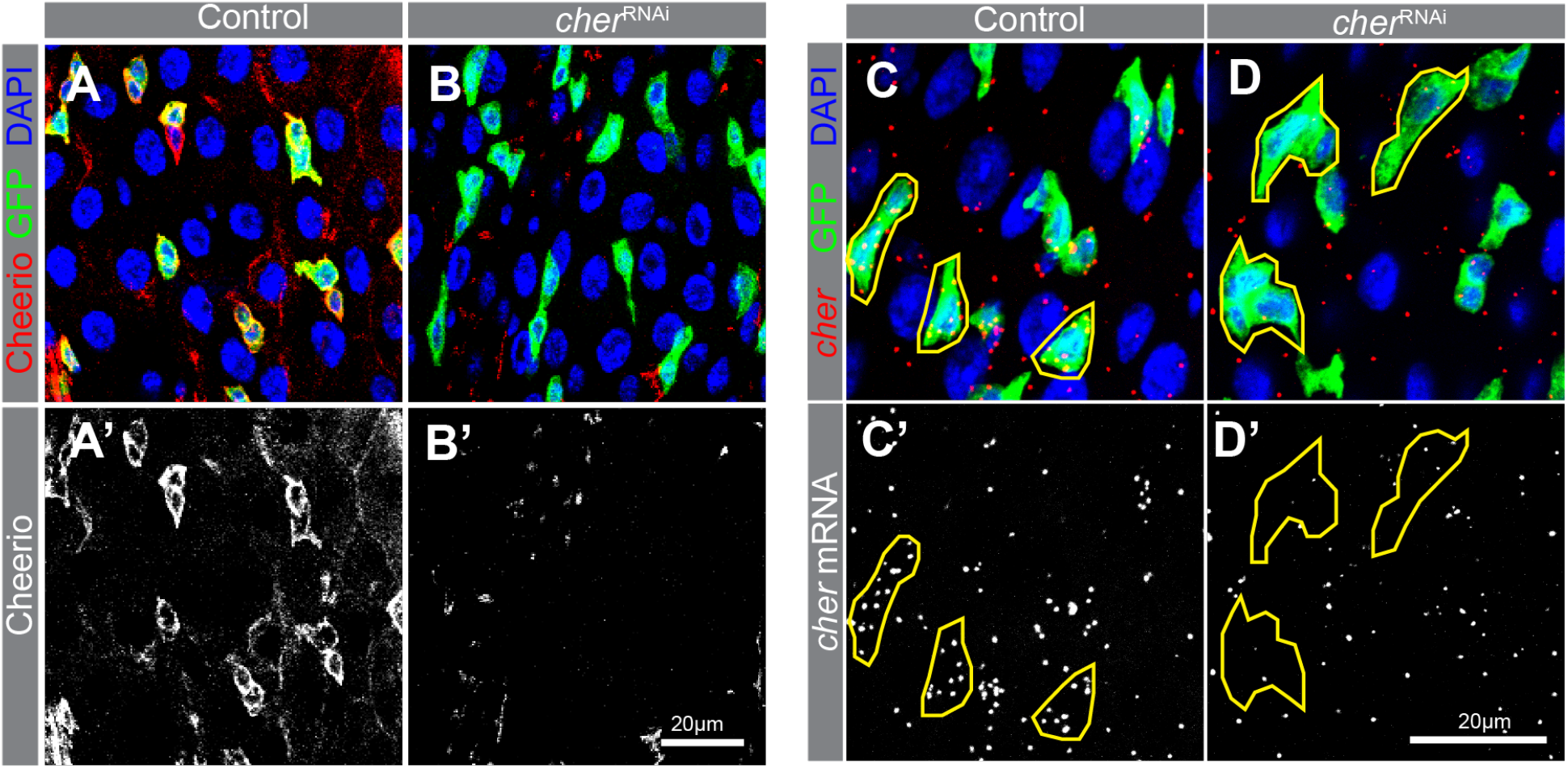
Verification of the *cher*^RNAi^ and *cher* RNAscope *in situ* probes. (A-B’) PMG sections from (A) *esg*^TS^ and (B) *esg*^TS^; *cher*^RNAi^ after 5 days at 29°C stained for Cheerio (anti-Cheerio, in red) and nuclei (DAPI, in blue). Ps are shown in GFP (green). (A’ and B’) Anti-Cheerio staining is shown in gray scale. (C-D’) Sections of PMG from *esg*^TS^ and (D) *esg*^TS^; *cher*^RNAi^ after 5 days at 29°C stained for *cher* mRNA (red) and Ps (anti-GFP, in green) and nuclei (DAPI, in blue). (C’ and D’) *cher* mRNA channel is shown in gray scale. Complete genotypes are listed in table S1. PMG, posterior midgut; P, progenitor cell.

